# Dynamics of the auditory continuity illusion

**DOI:** 10.1101/2021.03.05.433617

**Authors:** Qianyi Cao, Noah Parks, Joshua H. Goldwyn

**Author notes:** Correspondence: Joshua H. Goldwyn. These authors contributed equally to this work (Qianyi Cao, Noah Parks).

## Abstract

Illusions give intriguing insights into perceptual and neural dynamics. In the auditory continuity illusion, two brief tones separated by a silent gap may be heard as one continuous tone if a noise burst with appropriate characteristics fills the gap. This illusion probes the conditions under which listeners link related sounds across time and maintain perceptual continuity in the face of sudden changes in sound mixtures. Conceptual explanations of this illusion have been proposed, but its neural basis is still being investigated. In this work we provide a dynamical systems framework, grounded in principles of neural dynamics, to explain the continuity illusion. We construct an idealized firing rate model of a neural population and analyze the conditions under which firing rate responses persist during the interruption between the two tones. First, we show that sustained inputs and hysteresis dynamics (a mismatch between tone levels needed to activate and inactivate the population) can produce continuous responses. Second, we show that transient inputs and bistable dynamics (coexistence of two stable firing rate levels) can also produce continuous responses. Finally, we combine these input types together to obtain neural dynamics consistent with two requirements for the continuity illusion as articulated in a well-known theory of auditory scene analysis: sustained responses occur if noise provides sufficient evidence that the tone continues and if there is no evidence of discontinuities between the tones and noise. By grounding these notions in a quantitative model that incorporates elements of neural circuits (recurrent excitation, and mutual inhibition, specifically), we identify plausible mechanisms for the continuity illusion. Our findings can help guide future studies of neural correlate of this illusion and inform development of more biophysically-based models of the auditory continuity illusion.

## 1 INTRODUCTION

How do listeners in crowded and noisy environments create stable auditory streams in the face of interruptions and “background” noise? How do listeners identify the stops and starts of overlapping and interwoven sounds to correctly parse an auditory scene? Answering these questions is fundamental to understanding auditory perception and neural processing of sounds. A perceptual illusion that sheds light on dynamic processing of multiple sounds is the auditory continuity illusion [1] (also called temporal induction [2]). The continuity illusion can be elicited when noise interrupts a variety of sounds including tones, frequency glides, sentences [1, 2], and sound textures [3]. The common aspect of this illusion is that, when the noise is sufficiently loud and shares spectral content with the interrupted signal, listeners perceive a continuous, uninterrupted sound. This illusion reveals a tendency for the auditory system to maintain perceptual continuity when confronted with sudden changes in the auditory scene and to sustain perception of sounds that are present prior to some masking or distracting sound. The continuity illusion has been thoroughly studied since its discovery [4, 5].

The version of the continuity illusion we investigate is an interrupted tone that can be perceived as continuous if the interruption (a short interval in which no tone is presented) is filled with broadband noise, as depicted in Fig. 1. Results of listening experiments inform conceptual models for the illusion such as Bregman’s theories of auditory scene analysis [1]. Fundamental questions remain about the perceptual origin and neural basis for the illusion (viz. whether it is a cortical phenomenon [6, 7, 8] or created at subcortical stations [9]) and whether it is required that peripheral responses signal discontinuities in the sound [10]). Perhaps due to these uncertainties, there have been few efforts to model how the dynamic activity of neural populations can generate the continuity illusion. Previous works point toward several possible neural mechanisms that can give rise to the illusion: feedforward intra-cortical connections [6], nonlinear dynamic self-excitation [11], and short-term synaptic plasticity [12]). These few and disparate studies are motivation for further model-based investigations of the continuity illusion.

**Figure 1.**
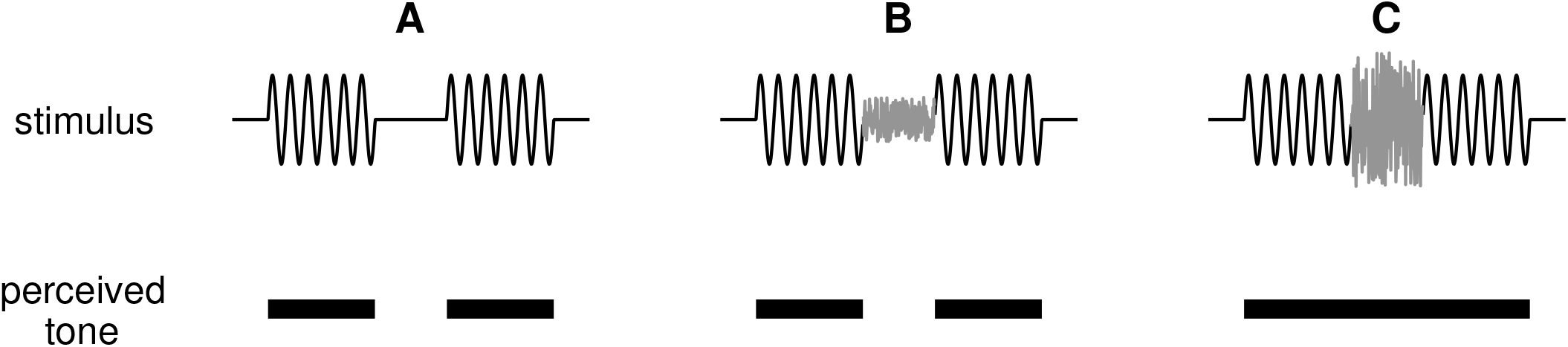
Illustration of the continuity illusion. (**A)** A tone interrupted by a silent gap (top panel) is perceived (correctly) as two tones discontinuous in time (bottom panel). (**B)** If a weak noise burst is inserted into the gap between tones, the tone remains perceived as discontinuous. (**C)** A sufficiently loud noise that shares frequency content with the tone can induce the illusion that the tone is uninterrupted and persists through the noise-filled interruption.

Our goal is to identify dynamical mechanisms that implement fundamental principles of the illusion. We are informed by the work of Petkov and colleagues who obtained evidence that two types of neural responses participate in the continuity illusion: sustained responses that signal ongoing sounds and transient responses that signal acoustic edges (onsets and offsets) [8]. While they specifically identified these response types in auditory cortex, we view these as common response motifs that are found throughout the auditory pathway. (see [13] for a review of offset auditory neurons, e.g.). In addition, we seek to connect Bregman’s principles, gained over decades of careful observation, to fundamental features of neural dynamics. Specifically, we present dynamical explanations for two of Bregman’s “rules” that define the circumstances under which the illusion will occur [1]. We paraphrase these rules as:

### Sufficiency of Evidence Rule

There must be some neural activity during the interruption that is indistinguishable from what would have occurred if the tone had continued through the noise-filled interruption.

### No Discontinuity Rule

There must be no evidence that the tone shuts off during the noise-filled interruption.

We use a nonlinear, dynamical firing rate model [14] with sustained inputs to implement the Sufficiency of Evidence rule. The dynamical principle at work is hysteresis: the interrupting noise (a broadband sound) provides a partial amount of excitatory input to an idealized neural population. Recurrent excitation in the population enables the noise to maintain firing activity in an already activated population, even though the noise alone cannot activate an inactive population. We then adjust excitability in the model so that it has two stable states that coexist in the absence of sustained inputs (bistability). We use this configuration to show that transient inputs at tone onsets and offsets implement the No Discontinuity rule. Continuity occurs when noise suppresses the offset response at the end of the first tone. Without this sufficiently strong offset response, the population remains active despite the absence of any ongoing input during the interruption between tones. Finally, we configure the model to receive both input types. In this setting, the model utilizes both hysteresis and bistability to create dynamics consistent with the continuity illusion. This formulation of using persistent neural activity as an indicator of perceptual state, instantiated as an attractor in a dynamical system, resembles ideas common to studies of the neural basis for working memory [15].

We illustrate our dynamical explanation for the continuity illusion with simulation result and we emphasize, throughout, analytical and geometric insights gained by computing equilibrium solutions to the nonlinear differential equation that governs the firing rate dynamics. By identifying principles of neural dynamics that are consistent with the continuity illusion, our work can guide future modeling studies that seek to simulate the illusion with more biophysically-detailed neural networks and inform studies that seek to identify the neural circuits and mechanisms that create the illusion.

## 2 MATERIAL AND METHODS

### 2.1 Firing rate model of a neural population

We model sound-evoked activity of a neural population using a firing rate equation with recurrent excitation [14]. The differential equation that describes the dynamics of the neural activity is

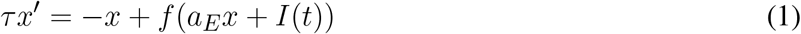

where *x*(*t*) is the firing rate variable that takes dimensionless values between 0 (population inactive) and 1 (population active). The value of *x* can be interpreted in a mean-field sense as the proportion of active neurons in a population or as the instantaneous probability of firing for neurons in the population. We suppose that an activity level near one for this population signals perception of a tone at some frequency. The parameter *τ* is a time constant of the firing rate dynamics and *a*_*E*_ is the strength of recurrent excitation within the population. The transfer function *f* is the sigmoidal nonlinearity

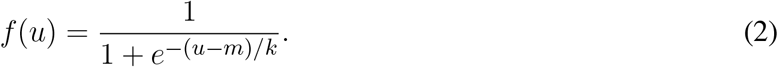

Since we are working with dimensionless quantities, we use *k* = 1 throughout without loss of generality. We explain our choices for the half-maximum parameter *m* in more detail below.

The input term *I*(*t*) is composed of for components:

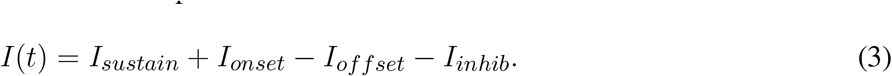

The first term is a sustained tone-driven excitatory responses (*I*_*sustain*_, constant for the duration of the tone). These second and third terms are transient, exponentially-decaying events triggered by tone onsets and offsets. The minus sign for *I*_*offset*_ indicates that offset responses are inhibitory inputs to the population. In some cases, there is also a noise-driven inhibitory term *I*_*inhib*_ that is a sustained response (constant for the duration of the noise). The classification of inputs as either sustained or transient, and the description of their dynamics as either constant or exponentially-decaying is a caricature of actual inputs motivated by observations by Petkov et al. [8]. Those authors proposed that sustained and transient response types in primary auditory cortex are possible neural correlates of the continuity illusion.

For responses to tones alone (no noise), the strengths of the tone-driven inputs are proportional to a dimensionless tone level parameter *I*_*T*_. By convention, we parameterize all models so that *I*_*T*_ = 1 is the threshold for tone perception (in the absence of noise). We also restrict *I*_*T*_ so it does not exceed a maximum tone level that we set (arbitrarily) at *I*_*T,max*_ = 5. The effect of noise varies by input type. Sustained inputs can be thought to represent the ongoing energy in the frequency-channel to which the population is tuned. Since noise is a broadband signal, we allow noise inputs to provide partial input to *I*_*sustain*_. Transient inputs can be thought to represent the salience of the onsets and offsets of a tone. We suppose noise degrades the “sharpness” of these acoustic edges and thus noise reduces *I*_*onset*_ and *I*_*offset*_ if noise is present at the instants of tone onset or offset. Finally, in some cases, we suppose there is a noise-driven pool that directly inhibits the tone population and that, in turn, receives inhibition from the tone population. For analytical convenience, we suppose this noise-driven inhibitory pool operates on a time-scale faster than the dynamics of *x*, and thus we let *I*_*inhib*_ evolve instantaneously to a value determined by noise strength. We use *I*_*N*_ to represent a dimensionless measure of noise level that takes value between 0 (threshold for noise perception) and *I*_*N,max*_ = 10 (arbitrarily chosen maximum noise level). From these considerations we have

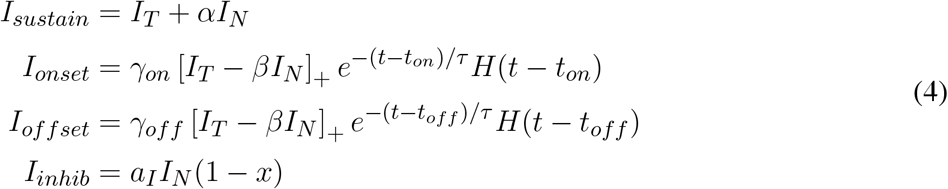

The parameters *α* and *β* are between 0 and 1 and represent the effects of noise on the tone-driven responses, as described above. The parameter *γ*_*on*_ scales the onset response so that tone threshold is always *I*_*T*_ = 1. We detail the calculation of *γ*_*on*_ below. The parameter *γ*_*off*_ similarly scales the offset response. We choose the decay time constant *τ* for transient responses to match the time scale of the firing rate dynamics. The notation [·]+ denotes rectification (values bound below at a floor of 0). The function *H* is the Heaviside function (also known as a step function) that is 0 if its argument is less than 0 and 1 if its argument is greater than 0. We use it here to indicate that the transient onset and offset responses begin at tone onset (*t*_*on*_) and tone offset (*t*_*off*_), respectively. The parameter *a*_*I*_ is the strength of the noise-driven inhibition. The multiplicative factor (1−*x*) in *I*_*inhib*_ implements mutual inhibition between the tone population and the noise-driven pool whose activity is represented by *I*_*inhib*_ since increases in the firing rate *x* reduce the strength of this noise-driven inhibitory input.

### 2.2 Stimulus configurations

We perform simulations and analyzing activation dynamics for three stimulus configurations: tone only, tone masked by noise, and tones interrupted by a noise-filled gap. In the masking configuration, tone and noise are presented simultaneously so that the noise impacts sustained and transient responses throughout the duration of the tone. In the interrupted tones configuration, the tone is shut off during the interruption between (*I*_*T*_ = 0) and noise impacts the offset response of the first tone and the onset response of the second tone.

As mentioned above, all models are parameterized so that *I*_*T*_ = 1 is the threshold for firing rate activation in the tone only case. In the masking case, we analyze activation dynamics and determine the smallest noise level at which a tone at a given level can overcome the effects of noise and activate the population. This defines the masking threshold *M* (*I*_*T*_). Similarly, for the interrupted tones, we determine the noise level (for a given tone level) at which the population remains active during the gap between tones. This defines the continuity threshold *C*(*I*_*T*_).

In simulations we use 1-second long inputs for tones (in tone only case) and tone and noise (in masking case). To simulate the continuity illusion, we present two tones at the same level and fill the interrupting gap with noise. The total duration of the continuity input is 2.5 seconds (tones are each 1 second long, the noise-filled gap is 0.5 seconds long). To be clear, tones and noise are represented by *I*_*T*_ and *I*_*N*_ in our model, we do not include sinusoidal waveforms for tones or stochastic processes for noise.

### 2.3 Numerical simulations

All calculations were carried out using the scientific computing software MATLAB (The MathWorks, Inc.). The firing rate differential equation (Eq. 1) was solved numerically using `ode15s`. Simulation code is available for download and use at https://github.com/jhgoldwyn/ContinuityIllusion. Nonlinear equations (to compute equilibrium states, for example) were solved using root-finding functions in Matlab such as `fzero` and `fsolve`.

## 3. RESULTS

### 3.1 Model classification by firing rate equilibria

We begin with a general analysis of the equilibrium states of the model for the case of sustained tone input only (*I*_*onset*_ = *I*_*offset*_ = 0 and *I*_*N*_ = 0). These calculations inform our parameter choices and are a starting point for subsequent analyses. Setting *x*′ = 0 and *I*(*t*) = *I*_*T*_ in Eq. 1) and solving for *I*_*T*_ we find the equilibrium relation

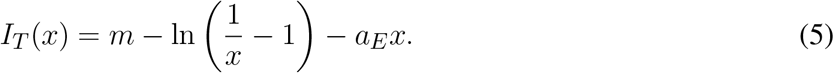

The parameters *a*_*E*_ and *m* determine the shape of *I*_*T*_ (*x*). We show three representative examples in Fig. 2A, with equilibrium firing rates plotted on the vertical axis and *I*_*T*_ plotted on the horizontal axis. The key features of these curves are: whether they are S-shaped with left and right “knees” and, if so, the values of *I*_*T*_ at these knees. Recall that the firing rate variable *x* takes values between zero and one. We interpret equilibrium solutions *x ≈* 0 as inactive states (no perception of tone) and equilibrium solutions *x ≈* 1 as active states (perception of tone).

**Figure 2.**
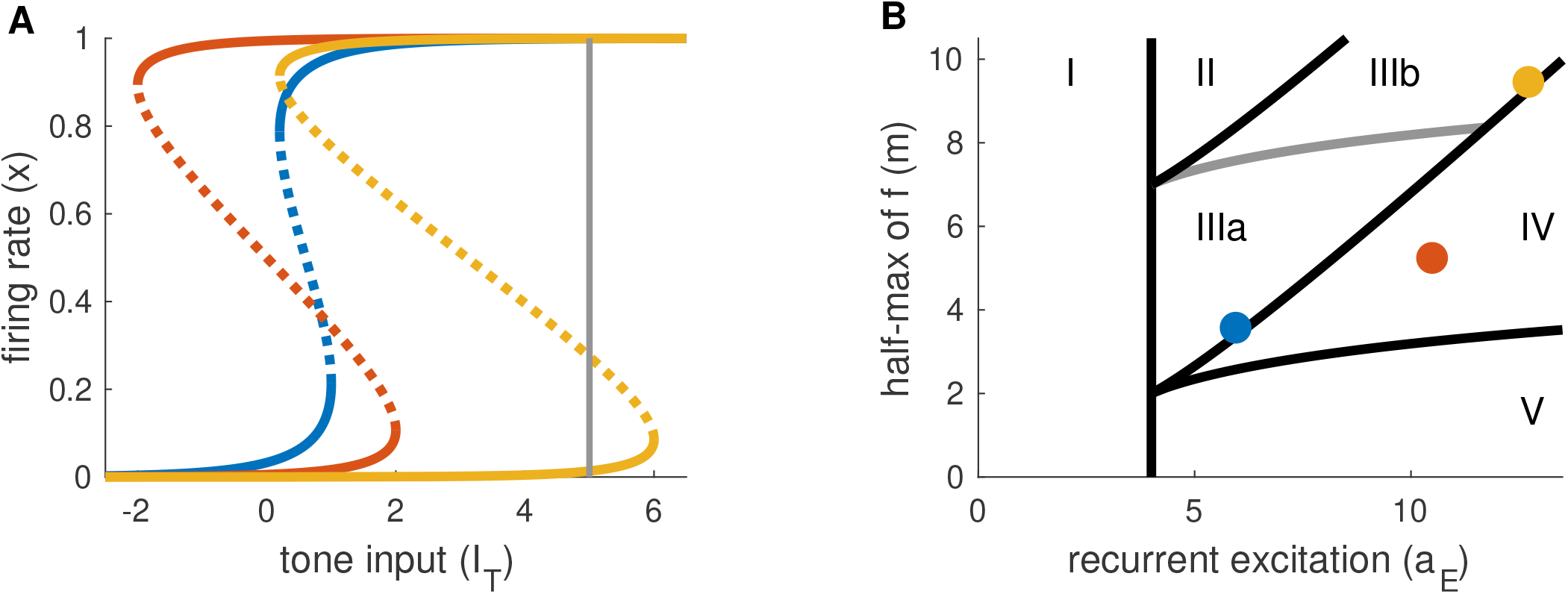
Partition of parameter space by shape of firing rate equilibrium curves. **(A)** Equilibrium firing rates for tone-only inputs (*I*_*N*_ = 0). Two stable branches (solid) are separated by a branch of unstable equilibria (dotted). Gray line at right indicates the maximum tone level used in simulations and analysis (*I*_*T,max*_ = 5). **(B)** Parameter space for parameters in Eqs. 1 and 2. Colored dots correspond to the curves in (A). Parameter space is partitioned according position of left and right knees of the firing rate equilibrium curves (see Table 1 for details).

We summarize the possible scenarios in Table 1 and show how they partition the *a*_*E*_-*m* plane in Fig. 2B. The scenarios that interest us must satisfy three criteria. First, to clearly distinguish between active and inactive states, the *x*-*I*_*T*_ curve must be S-shaped. This rules out Region I. Second, the left knee must be located at an *I*_*T*_ value less than the maximum tone strength (*I*_*T,max*_ = 5) so that activation is possible. This rules out Region II. Lastly, the right knee must be located at a positive *I*_*T*_ value so that inactivation is possible. This rules out Region V, in which the population could remain active at all times, even without inputs.

**Table 1.** Regions of *a*_*E*_-*m* parameter space classified by positions of left and right knees of *x*-*I*_*T*_ equilibrium firing rate curves. Regions of interest for this study are II and III. Other regions are ruled out because inactive and active states are indistinguishable (I), active state is inaccessible (II), or inactive state is inaccessible (V).

| Region | Left knee | Right knee |
| --- | --- | --- |
| I | none | none |
| II | $> I_{T,max}$ | $> I_{T,max}$ |
| IIIb | $(0, I_{T,max})$ | $> I_{T,max}$ |
| IIIa | $(0, I_{T,max})$ | $(0, I_{T,max})$ |
| IV | $< 0$ | $> 0$ |
| V | $< 0$ | $< 0$ |

As detailed in Table 1, the number and locations of the knees of the equilibrium curve delineate these regions. The knees are saddle node bifurcation points in the firing rate dynamics (points at which stable and unstable equilibria appear or disappear). They are located at critical points of *I*_*T*_ (*x*), so we identify these points by taking a derivative of Eq. 5 with respect to *x* and setting the resulting expression to 0. The *x* values at the left and right knees are

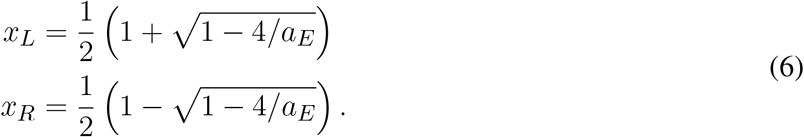

We compute the value of *I*_*T*_ at the knees by plugging these values into Eq. 5. In particular, we parameterize *a*_*E*_ and *m* by specifying *I*_*T*_ (*x*_*L*_) and *I*_*T*_ (*x*_*R*_) and solving the resulting two nonlinear equations given by Eq. 5. No real solutions exist for *a*_*E*_ *<* 4, thus *a*_*E*_ = 4 marks the boundary of models without S-shaped equilibrium curves (region I in Fig. 2B). The regions of interest for this study (III and IV) by the positions of the knees in the equilibrium curve. In particular, region IV consists of (*a*_*E*_, *m*) parameter pairs for which the model is bistable in the absence of any inputs (the left knee is at a negative *I*_*T*_ value and the right knee is at a positive *I*_*T*_ value). Region III consists of models that are monostable with no inputs (left knee is at a positive *I*_*T*_ value). We distinguish between two subregions of Region III. In Region IIIa, *I*_*T*_ at the right knee is less than the maximum tone value, and thus sustained inputs alone can activate the neural population. In contrast, in Region IIIb the *I*_*T*_ value at the right knee is larger than the maximum tone strength and thus models in this region can only be activated by a combination of sustained and transient inputs.

From these considerations, we choose three models (one from each region of interest) and use these in all further analysis and simulations.

**Model 1** (*a*_*E*_ = 5.9, *m* = 3.6, in Region IIIa). The left knee is at *I*_*T*_ = 0.2 and the right knee is at *I*_*T*_ = 1. As we describe below, the essential dynamical feature of this model is hysteresis: in response to sustained inputs the system requires a higher tone level to activate than deactivate (the S-shape of the equilibrium curve creates a mismatch between the two knees). We use sustained inputs only for this model (*I*_*onset*_ = *I*_*offset*_ = 0 in Eq. 3).

**Model 2** (*a*_*E*_ = 10.5, *m* = 5.2, in Region IV). The left knee is at *I*_*T*_ = −2 and the right knee is at *I*_*T*_ = 2. As we describe below, the essential dynamical feature of this model is bistability. We use transient inputs to move the system between active and inactive states and do not include sustained inputs (*I*_*sustain*_ = *I*_*inhib*_ = 0 in Eq. 3).

**Model 3** (*a*_*E*_ = 12.7, *m* = 9.5, in Region IIIb). The left knee is at *I*_*sust*_ = 0.2 and the right knee is at *I*_*sust*_ = 6. Sustained inputs alone cannot activate models in Region IIIb, so we use a combination of transient and sustained inputs for this model (all input terms in Eq 3 are non-zero).

Next, we show how each of these three models can demonstrate firing activity consistent with masking and the continuity illusion. We also compute masking and continuity thresholds analytically and explore effects of key parameters. We begin with Model 1 and show how it utilizes sustained inputs and the hysteresis effect to create firing rate dynamics consistent with the continuity illusion. We then show that Model 2, which uses only transient inputs but is set at a bistable point, can also produce the continuity illusion. Since sustained inputs signal (ongoing) evidence of a tone and transient inputs signal tone onsets and offsets, we view Model 1 as an implementation of Bregman’s No Discontinuity rule and Model 2 as an implementation of the No Discontinuity rule. Lastly, we show Model 3, which uses a combination of input types (sustained and transient), implements both rules for the illusion. Firing activity in Model 3 can persist through an interruption between tones because noise has two effects: to contribute excitatory inputs during the interruption (sufficient evidence), and to prevent the offset response from inactivating the population (no discontinuity).

### 3.2 Model 1: Sustained inputs and hysteresis dynamics implement the Sufficiency of Evidence Rule

#### 3.2.1 Response dynamics to tone-only inputs

Recall that Model 1 includes sustained inputs only (*I*_*onset*_ = *I*_*offset*_ = 0). The equilibrium solutions for this model for tone inputs only (*I*_*N*_ = 0) are shown in Fig. 3A1 with *I*_*T*_ as the bifurcation parameter (horizontal axis). When there is no tone input (*I*_*T*_ = 0), this system has a single stable equilibrium in the inactive state. For stronger tone inputs, the system passes through a saddle point bifurcation point at which a second stable equilibrium is created in the active state. Activation of the population from rest requires the tone to be larger, namely that *I*_*T*_ *> I*_*R*_ where *I*_*R*_ is the tone level at the right knee of the equilibrium curve. At this second saddle node bifurcation point, the inactive state is abolished and only the active state remains as the unique stable and globally attracting fixed point for the system. Responses to subthreshold and suprathreshold tone inputs are shown in Fig. 3A2.

**Figure 3.**
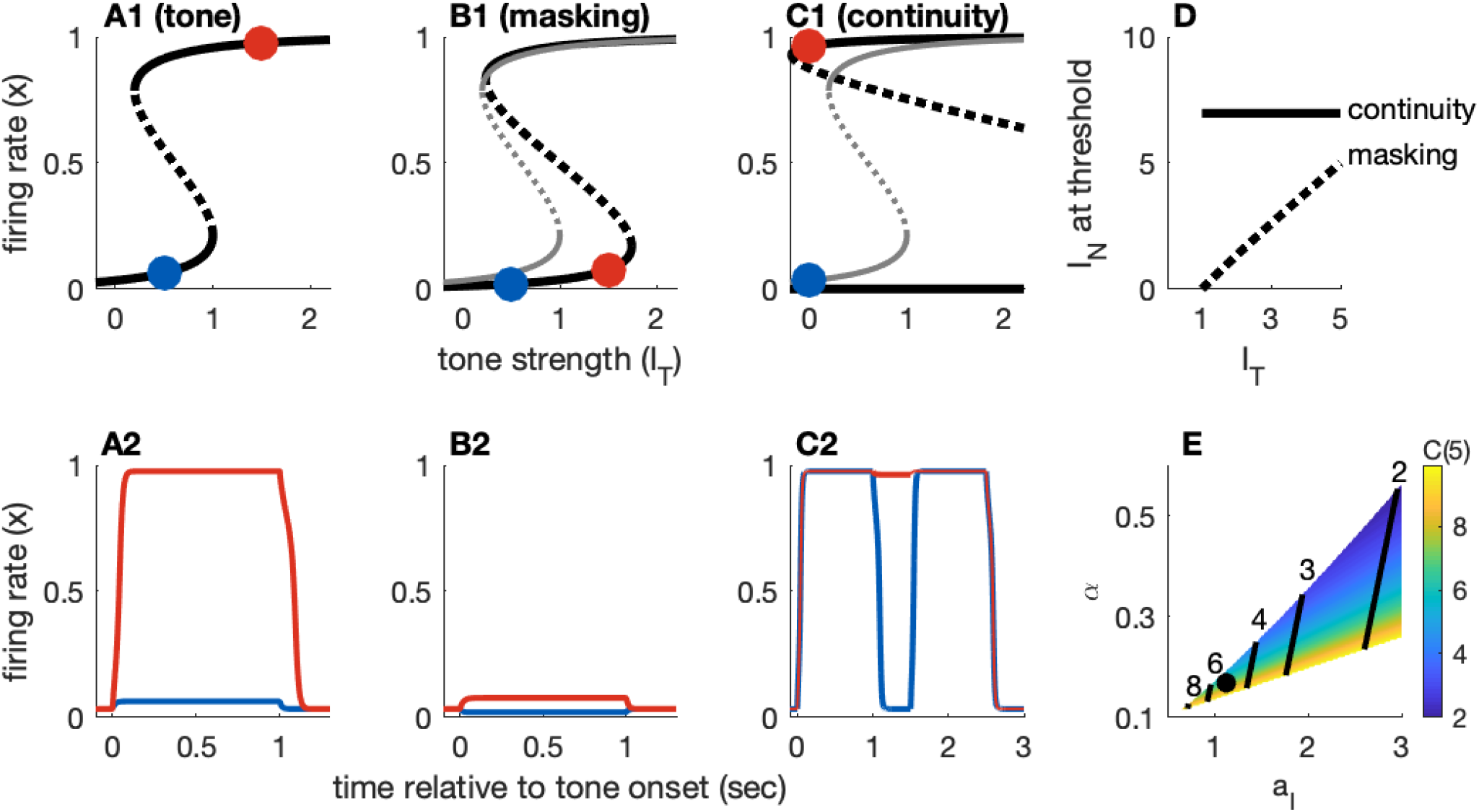
Dynamics of Model 1 responses to tone and noise inputs. **(A)** Tone inputs (no noise) with tone level *I*_*T*_ = 0.5 (blue) and *I*_*T*_ = 1.5 (red). Equilibrium firing rates shown in panel A1 and time-courses of firing rate variable *x*(*t*) in panel A2. Firing rate equilibrium in A1 (tone only) is re-plotted in gray in B1 and C1. **(B)** Simultaneous tone and noise inputs (masking condition) with *I*_*N*_ = 1. Noise shifts right knee of equilibrium curve to larger *I*_*T*_ levels (B1) and prevents activation by tone inputs (*I*_*T*_ = 0.5 and *I*_*T*_ = 1.5 shown). **(C)** Interrupted tone with noise-filled gap (continuity condition) with *I*_*T*_ = 1.5. Noise shifts left knee of equilibrium curve to smaller *I*_*T*_ levels (C1). Continuity occurs if left knee crosses *I*_*T*_ = 0 axis. Time-courses of firing rate variable in C2 for no noise (blue) and noise above continuity threshold (*I*_*N*_ = 8, red). (**D)** Masking threshold and continuity thresholds. (**E)** Masking and continuity thresholds at maximum tone level, for varying parameter *a*_*I*_ and *α* parameter values. *C*(*I*_*T,max*_) shown as a color map, contour lines show *M* (*I*_*T,max*_) (values labeled above contours). Parameter values used in panels (A - D): *a*_*I*_ = 1.124, *α* = 0.168, shown as black dot in E.

The feature of this model that is essential in our study of the continuity illusion (discussed below) is that it exhibits hysteresis dynamics. By this we mean that the tone level that activates an inactive population is larger than the tone level that maintains an already active population in the active state. Hysteresis is seen geometrically in the S-shaped *x*-*I*_*T*_ equilibrium curve (Fig. 3A1). The activation threshold is the tone strength at the right knee. The deactivation threshold – the minimum tone level that maintains activity – is the tone strength at the left knee. For Model 1, we positioned these knees at *I*_*T*_ = 1 (activation) and *I*_*T*_ = 0.2 (deactivation). As an example of the hysteresis effect: tone input with *I*_*T*_ = 0.5 does not activate *x* from rest (blue curve in Fig. 3A2) but it would maintain *x* at a level near 1 if *x* were active prior to this input (not shown, but notice the upper branch of equilibria in Fig. 3A1 extends to *I*_*T*_ values less than 0.5).

#### 3.2.2 Response dynamics to tone masked by noise

The two effects of noise are that it provides a “partial” input that enters as the additive term *αI*_*N*_ in *I*_*sustain*_, and that it produces noise-driven inhibition through the term *I*_*inhib*_ = *a*_*I*_*I*_*N*_ (1 − *x*) (recall Eq. 4). The effect of the inhibitory term is to shift the right knee of the equilibrium curve to larger *I*_*T*_ levels (Fig. 3B1, compare black curve with noise to gray curve without noise). That is, the threshold for activation increases with *I*_*N*_, as desired for noise to have a masking effect. As a demonstration, a tone level of *I*_*T*_ = 1.5 would activate the population from rest in the absence of noise, but with *I*_*N*_ = 1 this input does not activate the population (red curve in Fig. 4B2).

**Figure 4.**
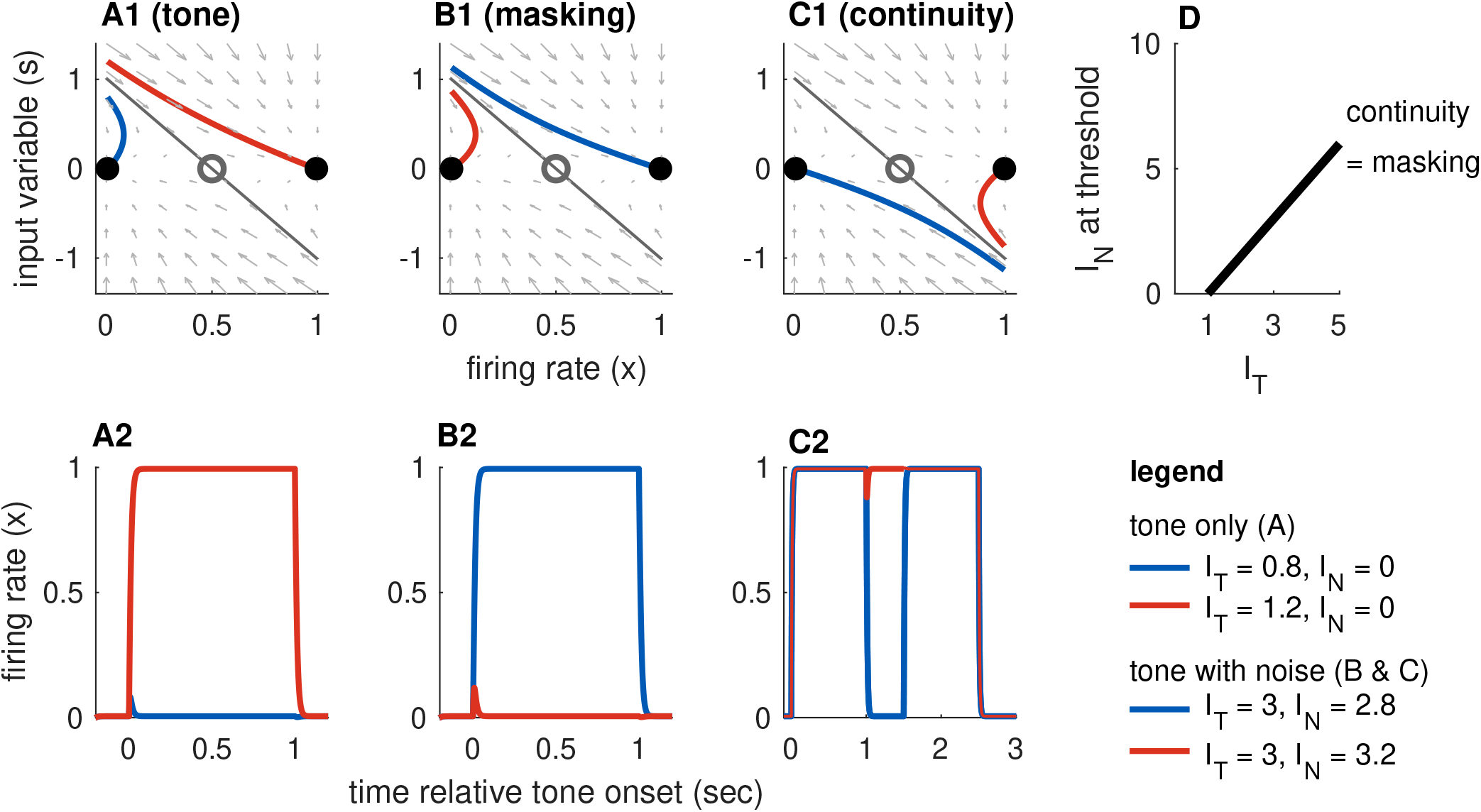
Dynamics of Model 2 responses to tone and noise inputs. **(A)** Tone inputs (no noise). Phase plane at tone onset, showing stable equilibria (black dots), unstable saddle (gray circle), linear approximation to separatrix (gray line), and trajectories for *I*_*T*_ = 0.8 (blue curve) and *I*_*T*_ = 1.2 (red curve). Time-course of firing rate variable *x*(*t*) shown in panel A2. **(B)** Simultaneous tone and noise inputs (masking condition) with *I*_*T*_ = 3. Phase plane shown at tone and noise onset. Sufficiently large noise suppresses the transient onset response and can prevent activation for sufficiently large noise (*I*_*N*_ = 3.2, red curve). **(C)** Interrupted tone with noise-filled gap (continuity condition) with *I*_*T*_ = 3. Phase plane shown at noise onset. Sufficiently large noise suppresses tone offset and can prevent return to inactive state (*I*_*N*_ = 3, red curve). (**D)** Masking threshold and continuity thresholds are equal for our choice of parameters (stable equilibria are equidistant to unstable saddle). Tone and noise levels shown in legend at bottom right. Parameter values used: *β* = 2*/*3, *γ*_*on*_ = *γ*_*off*_ = 5.2.

The equilibrium solutions for this model, using *I*_*T*_ as the bifurcation parameter and now including the effect of noise, are

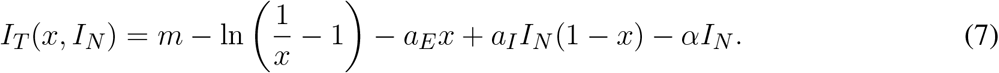

The critical points at which stable equilibria are created and abolished are the knees of the S-shaped curve. We obtain results similar to Eq. 6, but now with noise included:

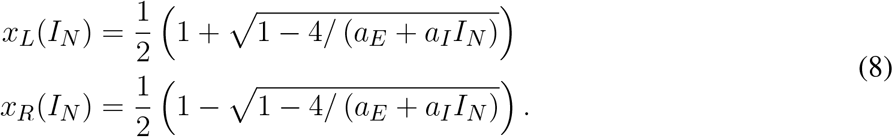

The *x*_*R*_(*I*_*N*_) point locates the threshold at which a tone activates a population from rest, in the presence of noise. The threshold for masking, then, is the noise level that solves *I*_*T*_ = *I*_*R*_(*I*_*N*_). Any value of *I*_*T*_ smaller than this critical value would fail to activate the population due to the noise-driven inhibition. We denote this masking threshold tone level as *M* (*I*_*N*_). It depends nonlinearly on *I*_*N*_ because of the dependence of *x*_*R*_ on *I*_*N*_. For Model 1, however, *x*_*R*_ is relatively constant with respect to *I*_*N*_ and we find masking threshold as a function of *I*_*T*_ is approximately linear (gray curve in Fig. 3D). The slope of this nearly linear relation can be approximated by [*a*_*I*_ (1 − *x*_*R*_(0)) − *α*]^−1^ (found by taking the derivative of *I*_*T*_ with respect to *I*_*N*_, and neglecting any change in *x*_*R*_ with respect to *I*_*N*_). This approximation shows the opposing effects of noise on masking threshold: *M* (*I*_*N*_) increases with increasing *α* (the amount of excitatory noise input) and decreases with increasing *a*_*I*_ (the amount of inhibitory noise-driven input), see Fig. 3H. This relation also imposes a constraint on model parameters. We must have *α < a*_*I*_ (1 − *x*_*R*_(0)). If this condition is not satisfied, the excitatory effect of noise (the *αI*_*N*_ term in *I*_*sustain*_) would dominate the inhibitory effect of noise (*I*_*inhib*_) and masking would not be possible.

#### 3.2.3 Response dynamics to tones interrupted by noise

The *x*_*L*_ point locates the threshold for inactivation in the presence of noise. A population in the active state will remain active during the gap between tones if the noise strength causes *I*_*T*_ (*x*_*L*_) to cross over to negative values, see Fig. 3C1. Thus, we calculate the continuity threshold equation by solving (numerically) the root-finding problem *I*_*T*_ (*x*_*L*_, *I*_*N*_) = 0, where *x*_*L*_ is also a function of noise level (Eq. 8). Observe that, since *I*_*T*_ = 0 during the gap between tones, the continuity threshold is constant with respect to tone level, as shown by the horizontal black line in Fig. 3D.

The masking and continuity thresholds for this model are separate. This imposes additional constraints on our parameter choices. Specifically, in accordance with the hypothesis that the continuity illusion is a compensation for masking [2, 16], we require that continuity can only occur at noise levels at least as high as the masking threshold. Additionally, we are only interested in parameter sets for which masking and continuity can both be achieved (continuity and masking threshold must not exceed *I*_*N,max*_ = 10, even for the strongest tones (*I*_*T,max*_ = 5). A view of the *a*_*I*_ − *α* parameter region that satisfies these requirements is shown in Fig. 3H, with labeled contour lines indicating the corresponding masking and continuity thresholds at the maximum tone level.

### 3.3 Model 2: Transient inputs and bistable dynamics implement the No Discontinuity Rule

3.3.1 Response dynamics to tone-only inputs

Model 2 receives transient inputs only (*I*_*sustain*_ = *I*_*inhib*_ = 0). These transient inputs occur at tone onsets and offsets and signal discontinuities at the “edges” of a tone. To analyze dynamics of this model at tone onset we find it useful to formulate it as a system of two ordinary differential equations:

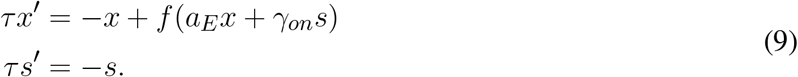

The additional variable *s* describes the exponentially-decaying transient input. At tone onset, this variable is instantaneously displaced to *s*(*t*_*on*_) = [*I*_*T*_ − *βI*_*N*_]+. Similar dynamics occur at tone offset, with *γ*_*on*_ replaced by *γ*_*off*_ and *s*(*t*_*off*_) = *—* [*I*_*T*_ − *β*_*I*_*N*]+. Notice *s*(*t*_*off*_) is negative-valued because offset responses are inhibitory inputs to *x*.

We configured Model 2 so that it is bistable in the absence of any inputs. In the *x*-*s* phase plane, bistability comprises stable equilibria at inactive and active firing rates (*x*_*I*_ and *x*_*A*_, respectively) separated by an unstable saddle point (*x*_*S*_). All equilibria are located along the *s*-axis. Activation of the population from rest requires that the transient onset response is sufficiently large to transition *x* from the basin of attraction of *x*_*I*_ to the basin of attraction of *x*_*A*_. This condition can be visualized in the phase plane by considering the separatrix curve *S*(*x*) that divides these two basins of attraction. The firing rate will activate from rest if *s*(*t*_*on*_) > *S*(*x*_*I*_), that is if the response variable at tone onset exceeds the height of the separatrix curve evaluated at the inactive firing rate equilibrium. We observed that the separatrix curve can be adequately approximated as a line connecting the saddle point (*x*_*S*_, 0) to the point (*x*_*I*_, 1). The choice of *s* = 1 at threshold enforces our convention that activation for tone-only inputs occurs at *I*_*T*_ = 1. From this geometric argument, the linear approximation to the separatrix is

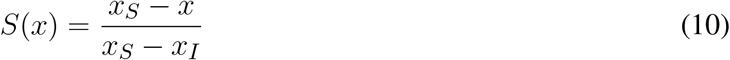

Trajectories in the phase plane at tone onset are shown in Fig. 4A1 and their full time-courses are shown in Fig. 4A2. If the tone level is sufficiently high (*I*_*T*_ = 1.2 in this simulation, red curve), the system crosses the separatrix and transitions to the stable equilibrium in the active state.

This approximation to the separatrix can also be found by linearizing the dynamical system in Eq. 9 about the saddle point and determining the eigenvector associated with its stable manifold. The Jacobian matrix for the system is

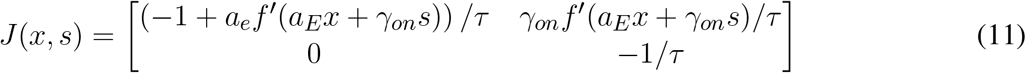

The negative eigenvalue for the saddle point is the lower right entry of this matrix. The associated eigenvector satisfies (*J*_11_ − *J*_22_)*x* + *J*_12_*s* = 0, where *J*_*ij*_ is the (*i, j*) entry of the Jacobian matrix so we conclude that the linear approximation to the separatrix is

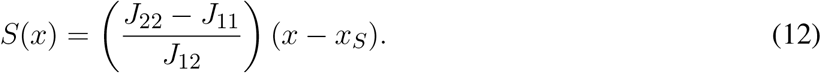

A useful consequence of our assumption that *x* and *s* have the same time constant *τ* is that this expression can be simplified substantially to:

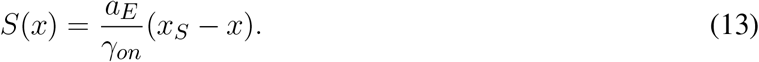

Comparing this equation to the result obtained by geometric considerations (Eq. 10), we observe that *γ*_*on*_ is determined by the parameters *a*_*E*_ and *m*, and our convention that tone threshold is *I*_*T*_ = 1. In particular, we set *γ*_*on*_ = *a*_*E*_(*x*_*S*_ − *x*_*I*_).

Deactivation is the mirror image of activation and occurs if the “downward” perturbation of *s* is sufficiently strong at the tone offset. For the values of *a*_*E*_ and *m* that we use, the stable equilibria are symmetric around the saddle point at *x* = 0.5 and it is convenient to set *γ*_*on*_ = *γ*_*off*_ so that the threshold for activation and deactivation are the same. More generally, the offset parameter should always be set so that activation thresholds are not less than deactivation thresholds, to avoid the scenarios in which a tone onset can activate the firing rate variable but the tone offset response is too weak to return the population. In this unrealistic case, *x* could remain in the active state for perpetuity.

#### 3.2.2 Response dynamics to tone masked by noise

If a sufficiently strong noise is presented at the same time as a tone, then the noise can prevent activation of the tone population by reducing the transient response at the start of the tone. This is the masking condition. Simulations exhibiting masking dynamics are in Fig. 4B. Recall the effect of noise is to reduce the onset response to *s*(*t*_*on*_) = [*I*_*T*_ − *βI*_*N*_]+ where *β* is a parameter that controls how much noise suppresses the tone onset (given in Eq. 4). As we explained above, in our discussion of tone activation, the model is parameterized so that *I*_*T*_ = 1 is the threshold for activation in the absence of noise. Thus, a noise will mask a tone if *I*_*T*_ − *βI*_*N*_ *<* 1. We denote the threshold for masking as *M* (*I*_*T*_) and conclude that it is related to tone strength via a linear equation:

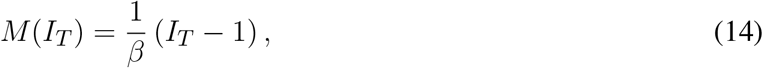

This equation is valid for values of *I*_*T*_ above the noise-free threshold (*I*_*T*_ = 1) and below the maximum tone strength in the model (*I*_*T,max*_ = 5). It provides a direct relationship between masking threshold and the degree to which noise masks tone onsets (represented by the parameter *β*). In the simulations shown in Fig. 4) we use *β* = 2*/*3. The masking threshold curve is shown in Fig. 4D.

#### 3.3.3 Response dynamics to tones interrupted by noise

The feature of this model that is essential for our study of the continuity illusion is that the inactive and active coexist and are stable in the absence of any inputs. The tone population can, therefore, remain in the active state even after a tone is turned off if the offset signal is weak and does not send *x* across the separatrix. Simulations exhibiting masking dynamics are in Fig. 4C.

Whereas masking depends on suppression of the onset response (as described above), the continuity illusion depends on suppression of the offset response. Computation of the continuity threshold is analogous to our derivation of the masking threshold, but with onset terms replaced by offset terms. The criteria for continuity are that *I*_*T*_ > 1 (so that the first tone activates the population) and that *I*_*N*_ is sufficiently large to reduce the tone offset response and allow the *x* firing rate to remain near the upper equilibrium state. To satisfy this second condition, we must have that the offset response does not cross the separatrix curve, with the key difference being that we are now analyzing the system at the start of the noise-filled gap. This means *x*(*t*_*off*_) = *x*_*A*_ (first tone has activated the population) and *s*(*t*_*off*_) = − [*I*_*T*_ − *βI*_*N*_]+. Adapting the separatrix equation in Eq. 13 for the offset response, we have that continuity requires – [*I*_*T*_ − *βI*_*N*_]+ < *a*_*E*_ (*x*_*S*_ − *x*_*A*_) */γ*_*off*_. The continuity threshold equation is, therefore,

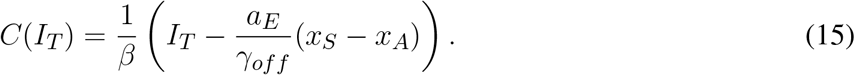

In the particular case of Model 2, we have chosen parameters that make it symmetric (*x*_*I*_ and *x*_*A*_ are equidistant from the saddle point). We set *γ*_*off*_ = *γ*_*on*_ = *a*_*E*_(*x*_*S*_ − *x*_*I*_), so that the continuity and masking thresholds are identical (compare *C*(*I*_*T*_) to Eq. 14. Masking and continuity threshold lines are identical and plotted in Fig. 4D).

More generally, we would require *γ*_*off*_ ≥ *γ*_*on*_ to avoid persistent activation (discussed above). If *γ*_*off*_ > *γ*_*on*_, the continuity threshold would shift upward in Fig. 4D (*γ*_*off*_ affects the intercept of *C*(*I*_*T*_) but not its slope). In the intermediate *I*_*N*_ values between *C*(*I*_*T*_) and *M* (*I*_*T*_), we observe responses to interrupted tones in which the first tone activates *x*, the noise does not cause firing to persist (no continuity) and the noise prevents activation by the second tone. In other words, the model without symmetry results in a region in stimulus space in which the noise burst between the two tones is too weak to induce the continuity illusion but sufficiently strong to prevent perception of the second tone by forward masking.

### 3.4 Model 3: Combined inputs implement both rules for the continuity illusion

The last model configuration we consider is one that cannot be activated by transient inputs alone or sustained inputs alone. These requirements are met if *I*_*T*_ at the left knee of the *I*_*T*_ -*x* equilibrium curve is positive (no bistability at rest) and the *I*_*T*_ at the right knee is to the right of the maximum allowable tone level *I*_*T,max*_. Activation from rest can only occur using a combination of sustained and transient inputs. There must be a sufficiently strong sustained input to move the system past the saddle node bifurcation point at the left knee of the equilibrium curve. This creates a stable equilibrium in the activated state that can be accessed if the transient portion of the input is sufficiently strong to transition the system into the basin of attraction of this upper equilibrium, see Fig. 5.

**Figure 5.**
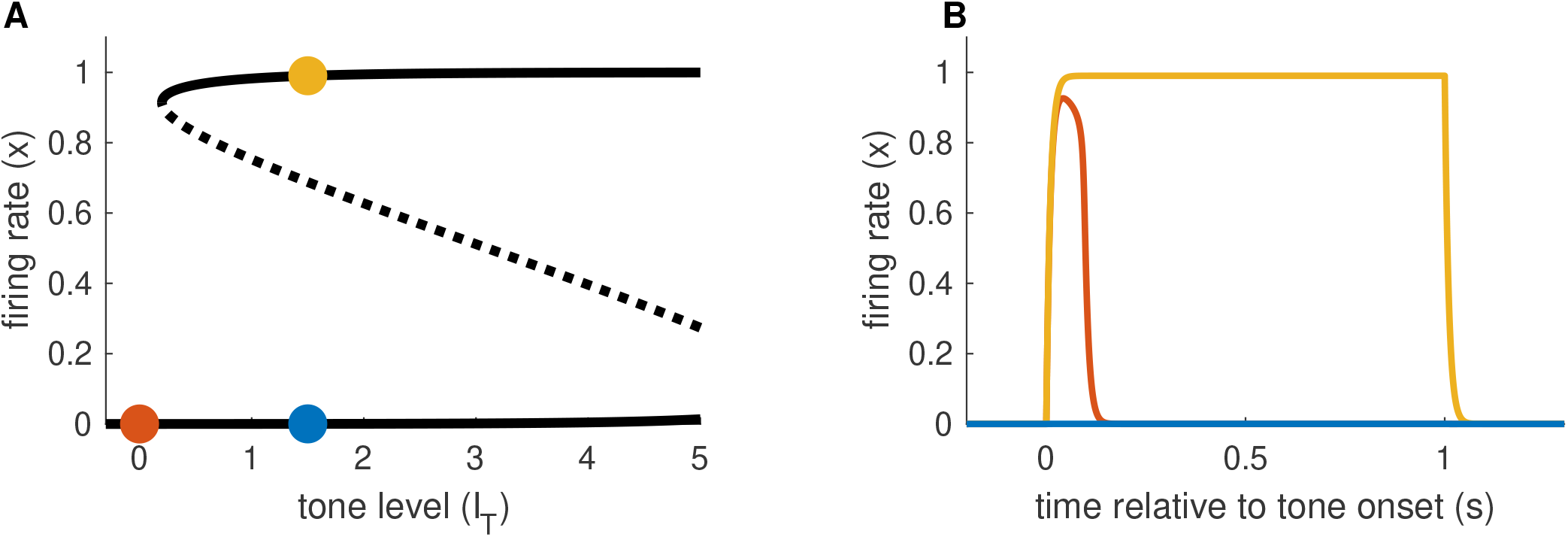
Activation dynamics in Model 3 requiring sustained and transient inputs. (**A)** Firing rate equilibrium curve. Defining feature of Model 3 is a left knee at positive *I*_*T*_ level and right knee at *I*_*T*_ level larger than *I*_*T,max*_. (**B)** Time-courses of firing rate variable for sustained input only (blue), transient input only (red), and combination of both inputs (yellow). Tone level is *I*_*T*_ = 1.5 in all cases.

To understanding firing rate responses for this model, we again formulate the dynamics in the *x*-*s* state space:

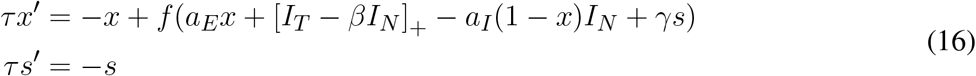

These equations govern the dynamics of the system in the time following a tone onset or offset. The parameter *γ* should be thought of representing *γ*_*on*_ to describe onset responses or *γ*_*off*_ for offset responses. The initial values for these equations are given by the state of the systems immediately prior to sound onset or offset. For the case of tone onset for a system starting from rest (no input), for instance, the initial values would be *x*(*t*_*on*_) = *x*_*I*_(0, 0) and *s*(*t*_*on*_) = 1, where *x*_*I*_(0, 0) is the inactive state in the case of *I*_*T*_ = *I*_*N*_ = 0. We will use similar notation throughout this section to indicate that equilibrium points are functions of the input levels.

The Jacobian matrix for these equations is

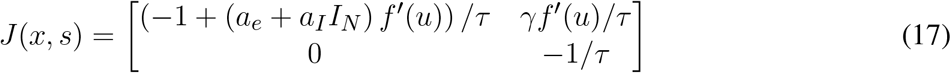

where we have abbreviated the argument of *f*′ with *u* = *a*_*E*_*x* + [*I*_*T*_ − *βI*_*N*_]+ − *a*_*I*_(1 − *x*)*I*_*N*_ + *γ*_*on*_*s*. As before, we evaluate the Jacobian at the saddle point (when it exists, for sufficiently large *I*_*T*_) and use the eigenvector associated with the stable manifold of the saddle point to construct a linear approximation to the separatrix curve that defines the threshold for activation. The notable difference between the analysis in this section and the preceding section (for Model 2), is that the sustained input terms can affect the positions of the equilibrium solutions and the shape of the separatrix in the current model setting. Following our earlier calculation (recall Eq. 12), we find the eigenvector by solving (*J*_11_ − *J*_22_)*x* + *J*_12_*s* = 0. After simplifications, we find the linear approximation to the separatrix to be

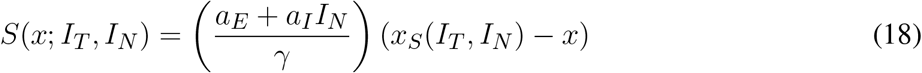

where we are assuming that *I*_*T*_ is sufficiently large so that the saddle point *x*_*S*_(*I*_*T*_, *I*_*N*_) exists. In the remaining sections, we apply this result in the three cases we have been considering (tone only, simultaneous tone and noise, and tones with a noise-filled gap) to characterize activation by tone, masking, and continuity dynamics.

#### 3.4.1 Response dynamics to tone-only inputs

Activation by a tone-only input (*I*_*N*_ = 0) occurs if the onset response causes the system to cross the separatrix defined in Eq. 18. In particular, we consider the system starting from rest, with *x*(*t*_*on*_) = *x*_*I*_(0, 0) and input variable *s* instantaneously perturbed to *s*(*t*_*on*_) = *I*_*T*_. We then ask if this onset perturbation to *s* exceeds *S*(*x*(*t*_*on*_)). We must also keep in mind that the position of the saddle point is determined by inputs, so in this case we use *x*_*S*_(*I*_*T*_, 0) in Eq. 18. From these considerations we conclude that, to satisfy our convention that *I*_*T*_ = 1 is the tone threshold, we must set the onset parameter to *γ*_*on*_ = *a*_*E*_ (*x*_*S*_(1, 0) − *x*_*I*_(0, 0)). Simulations showing activation by a tone are shown in Fig. 6A. The phase portrait in Fig. 6A1 illustrates the dynamics at tone onset. We remark that *I*_*offset*_ is not necessary to move the system back to the inactive state. In this model, the return to the inactive state at the end of the tone is guaranteed in the tone-only case because the saddle point and upper equilibrium do not exist for *I*_*T*_ = 0. The firing rate variable must return to *x*_*I*_(0, 0) because it is the unique, remaining stable equilibrium. This differs from Model 2 which required an offset response to deactivate *x*. We will see shortly, however, that the offset response does affect dynamics of the continuity illusion dynamics, and we will explore *γ*_*off*_ further in that setting.

**Figure 6.**
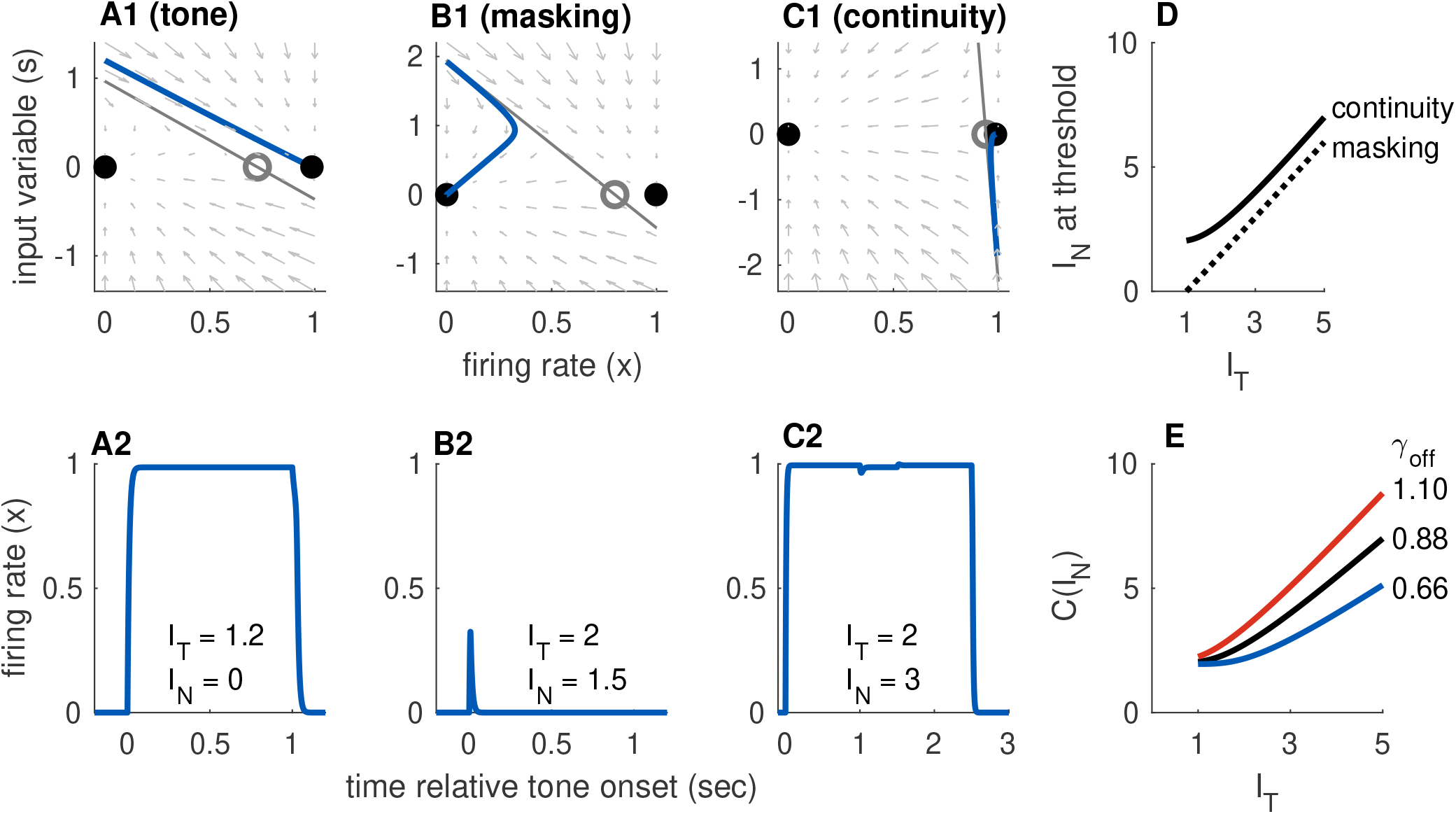
Dynamics of Model 3 responses to tone and noise inputs. **(A)** Tone inputs (no noise) with tone level *I*_*T*_ = 1.2. Phase plane at tone onset, showing stable equilibria (black dots), unstable saddle (gray circle), linear approximation to separatrix (gray line) and trajectory (blue curve). Time-course of firing rate variable *x*(*t*) shown in panel A2. **(B)** Simultaneous tone and noise inputs (masking condition) with *I*_*T*_ = 2 and *I*_*N*_ = 1.5. Phase plane shown at tone and noise onset. The active state still exists, but it is shifted rightward relative to tone only and slope of the separatrix is steeper. Noise prevents activation in this case and firing rate returns to inactive equilibrium. **(C)** Interrupted tone with noise-filled gap (continuity condition) with *I*_*T*_ = 2 and *I*_*N*_ = 1.5. Phase plane shown at noise onset. Noise preserves stable equilibrium. In this case, trajectory does not cross separatrix and system remains active during gap between tones. (**D)** Masking threshold and continuity thresholds. Parameter values used: *a*_*I*_ = 7, *α* = 0.5, *β* = 0.05, *γ*_*on*_ = 9.6, *γ*_*off*_ = 0.88. (**E)** Continuity threshold for varying *γ*_*off*_ parameter value. Black curve in (D) and (E) are identical.

#### 3.4.2 Response dynamics to tone masked by noise

In the masking condition, tone and noise inputs are both present simultaneously. As a result, the saddle point about which we linearize the system depends on the inputs *x*_*S*_(*I*_*T*_, *I*_*N*_). To determine whether a noise prevents activation by a tone, we are still interested in onset dynamics from rest. The initial firing rate is *x*(*t*_*on*_) = *x*_*I*_(0, 0). We use the separatrix equation (Eq. 18) again to determine the masking threshold. We determined *γ*_*on*_ and we use it to update the separatrix at tone onset:

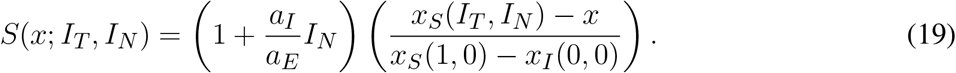

This equation reveals the differing effects of sustained tone and noise inputs. The sustained tone input does not alter the slope of the linear approximation to the separatrix, it is the same as the slope found for the model with transient inputs only (Eq. 12), but it can change the intercept through the dependence of *x*_*S*_ on *I*_*T*_ in the numerator. The sustained noise input, in contrast, changes the intercept and also the slope of the intercept (through the term *a*_*I*_ *I*_*N*_ */a*_*E*_).

Trajectories in the phase plane and as firing rate time-courses showing masking of a tone by noise are in Fig. 6B. In this example, the onset response exceeds the threshold for tone-only inputs (*s*(*t*_*on*_) is above one in Fig. 6B1). Nevertheless, the population returns to the inactive state because the noise input has raised the threshold for activation (in this case, primarily by steepening the slope of separatrix).

With Eq. 19 in hand, we compute the masking threshold as the minimum *I*_*N*_ level (for a given *I*_*T*_ level) at which the onset response crosses the separatrix. To do this, we set the amplitude of the transient onset response to the height of the separatrix at the initial value. Thus, the masking threshold *M* (*I*_*T*_) is the noise level that solves the nonlinear equation *I*_*T*_ − *βI*_*N*_ = *S*(*x*_*I*_ (0, 0); *I*_*T*_, *I*_*N*_). We solve this equation numerically and display the resulting masking threshold curve in Fig. 6D, for selected parameter values.

#### 3.4.3 Response dynamics to tones interrupted by noise

In contrast to the two cases just considered (activation by a tone alone, masking by noise), understanding the continuity illusion requires analysis of the offset response. We approximate the threshold, as usual, with the linear approximation to the separatrix that is given in Eq. 18. We interpret *γ* in that equation as *γ*_*off*_ since we are concerned with the dynamics at the offset of the first portion of the tone. There is no tone (*I*_*T*_ = 0) during the noise-filled interruption between the tones, so the saddle point position is only a function of *I*_*N*_. This highlights the fact that there are two necessary conditions for continuity dynamics in this model: the noise level must be sufficiently strong to preserve the saddle point and stable active equilibrium and the offset response must be sufficiently weak (perhaps because it is masked by the noise) to prevent the system from crossing the separatrix. The second of these conditions is satisfied if the offset response, which has initial amplitude – (*I*_*T*_ − *βI*_*N*_), does not cross *S* (*x*_*A*_; 0, *I*_*N*_). A firing rate response that remains active when the tone is removed (consistent with the continuity illusion) is shown in the *x*-*s* phase plane in Fig. 6C1 and as a time-course in Fig. 6C2.

The offset parameter *γ*_*off*_ shifts the system between the two extreme cases represented by Models 1 and 2. If the offset parameter is small, then the occurence of the continuity illusions relies on the hysteresis effect (the fact that *I*_*N*_ can preserve the saddle point and active equilibrium). In this case, the continuity threshold *C*(*I*_*T*_) changes slowly with *I*_*T*_ (and, in fact, can be constant over a range of *I*_*T*_ values). In the extreme case of no offset response, the continuity threshold would be constant with *I*_*T*_ as was the case for Model 1 (see Fig. 3D). If the offset parameter is large, then the offset response makes a larger contribution to whether the firing persists during the gap. The continuity threshold varies more with *I*_*T*_ as *γ*_*off*_ increases. These effects of *γ*_*off*_ on continuity threshold are shown in Fig. 4D.

## 4 DISCUSSION

The continuity illusion is an intriguing example of the capacity for the brain to “fill in” missing information. In this case, this “filling in” process creates an illusion of a continuous tone that is, in fact, discontinuous. In the context of hearing in a complex listening environment, a bias toward linking related sounds across time to create longer-lasting auditory objects can be useful when contending with multiple interrupting or distracting sounds that could momentarily obscure a sound of interest. Our contribution has been to use principles of neural dynamics and to draw on previous conceptual frameworks and experimental studies of the continuity illusion to identify dynamical mechanisms that can account for the continuity illusion. Our approach was to use a firing rate model as a caricature of population-level neural activity. Firing rate models have been used in studies of visual [17, 18, 19] and auditory illusions [20, 21, 22] and are useful for identifying possible dynamical and neural mechanisms that underlie perceptual outcomes.

We explored the possible routes toward continuity illusion-like dynamics. A first requirement was that recurrent excitation be strong enough to create a stable equilibrium at a high firing rate level (Fig. 2). We stereotyped the inputs to the population as sustained, transient onset, and transient offset. This setup was based on physiological evidence that neurons in auditory cortex with these response types are possible neural correlates for the continuity illusion [8]. Although the model could be configured so that sustained inputs alone (Model 1) or transient inputs alone (Model 2) produce dynamics consistent with the continuity illusion, there are shortcomings in both cases that can be remedied in a model that requires both sustained and transient inputs together to activate the population (Model 3). In the case of sustained inputs alone, continuity dynamics require a hysteresis effect: input levels that are too weak to activate the population from rest can, nevertheless, sustain activity in an already active population. The tone input makes no contribution to the response dynamics during the noise-filled interruption between the tones. As a result, the continuity threshold is constant with tone level (Fig. 3D). This is inconsistent with evidence that the probability of perceiving the continuity illusion increases as the noise level becomes louder relative to the tone [23]. In the case of transient inputs alone, continuity dynamics require a population that is bistable in the absence of any inputs. If additional mechanisms were not at work (synaptic adaption in the recurrent excitatory connections, for instance), a bistable population could remain in a high firing rate state for a long period of time even when no stimulus is present. The model that requires both inputs to activate (Model 3) can utilize both input types to implement the continuity illusion. Sustained noise-driven inputs are necessary to preserve a stable equilibrium in the active state (via a hysteresis effect) and transient offset responses can ensure that the continuity threshold is not independent of tone level (Fig. 6H).

Following Petkov et al. [8], we view the different input populations (sustained, onset, and offset) as possible neural correlates of two aspects of the descriptive theory of the continuity illusion proposed by Bregman [1]. First, sustained responses convey “evidence” that a tone is ongoing. If some sustained neural activity persists during the interruption between two tones, then the tone may be perceived as continuous. This is Bregman’s “Sufficiency of Evidence rule.” In our model, this is implemented by assuming that noise (as a broadband signal) drives a partial excitatory input to the population whose activity signals the perception of the tone. Second, transient responses at tone onsets and offsets mark “discontinuities” in the tone (acoustic edges). If the noise during the gap between tones obscures this discontinuity, then tone may be perceived as continuous. This is Bregman’s “No Discontinuity rule.” In our model, this is implemented by assuming noise can reduce the amplitude of the transient response. An additional effect of noise (in Model 3) is to increase the threshold for activation by a transient input (effected by altering the separatrix, see *I*_*N*_ term in Eq. 19).

By analyzing how these input types drive firing rate activity, we identified the dynamical mechanisms and circuit properties that can support the continuity illusion. A hysteresis effect enabled the noise to convey “sufficient evidence” to sustain firing activity in the gap between tones. A suppression of the offset response due to noise kept removed the “discontinuity” in the acoustic signal and confined the system to the basin of attraction of the high firing rate stable equilibrium. These dynamical mechanisms (hysteresis and bistability) required sufficiently strong recurrent excitation, specifically the requirement that *a*_*E*_ > 4 for S-shaped equilibrium curves (Fig. 2).

In addition to providing a dynamical framework to accompany Bregman’s theory, our approach offers a new perspective on previous models of the continuity illusion. The work of Noto et al. [11] relied on nonlinear, self-excitation of neural populations which is consistent with our approach for creating hysteresis and bistable dynamics. The work of Vinnik et al. [12] utilized dynamic synapses to create continuous responses to interrupted tones. Since dynamic synapses are an additional mechanism for creating hysteresis and bistable dynamics [24], there is also consistency between their work and ours. An extra feature of our study, not considered in these previous models of the continuity illusion, is that we also required that our simulated dynamics exhibit masking for noise and tones presented simultaneously. Incorporating this is essential, in our view, to avoid “trivial” solutions in which noise acts simply (and solely) as an excitatory input that facilitates neural activity during the noise-filled gap between tones. Balancing these excitatory and inhibitory effects of noise required fine-tuning in Model 1 (Fig. 3E).

Although the firing rate framework is a caricatured description of neural activity, insights into a number of sensory illusions have been made using similar approaches including for visual bistability [17, 18, 19] and auditory streaming [20, 21, 22]. In our study, it has been useful as a minimal model that identifies mechanisms that can account for the continuity illusion. We focused on a classic version of the continuity illusion (tone interrupted by noise), but the illusion can be elicited by a variety of sound types [1, 2, 3]. The facts that the continuity illusion is widespread and that the dynamics of the illusion can be explained with a relatively simple model may indicate that circuitry that promotes perceptual continuity is present at multiple auditory centers. Indeed, while we were motivated by observations in auditory cortex [8], the input motifs we used in our model (sustained, onset, and offset responders) are present in multiple auditory nuclei. See, for instance, Rhode and Smith [25] for sustained and onset response types in the ventral cochlear nucleus and Kopp-Scheinpflug et al. [13] for descriptions of offset neurons at multiple nuclei in the auditory system). There is also evidence for involvement of the brainstem in the continuity illusion in human listeners [9]. These observations may support a view that perceptual continuity of interrupted sounds is constructed at more than one stage of auditory processing. As work continues to identify neural correlates of the continuity illusion, idealized dynamical systems descriptions can inform what features should be included in future, more biophysically-detailed modeling approaches.

## RESOURCE IDENTIFICATION INITIATIVE

Computations were performed using MATLAB (RRID:SCR 001622), version R2018b.

## AUTHOR CONTRIBUTIONS

JHG conceived of the study. All authors designed, implemented, and carried out simulations and analysis. All authors contributed to writing and editing the manuscript.

## DATA AVAILABILITY STATEMENT

Matlab code for the firing rate model and to generate all figures in this study can be found in the github repository at https://github.com/jhgoldwyn/ContinuityIllusion.

